# Restoration of tropical dry evergreen forest in southern India: balancing carbon sequestration with biodiversity conservation

**DOI:** 10.64898/2026.07.08.737378

**Authors:** Shanmugam Mani, Sandeep Pulla, Lourde Nadin Epinal

## Abstract

Tropical dry evergreen forests (TDEFs) are a unique and highly threatened forest type of the dry tropics. Their restoration could be strengthened if native species demonstrate carbon sequestration comparable to widely used non-native trees. We assessed biodiversity and carbon sequestration in a restored TDEF in India, developed over 50 years from a largely barren landscape. The site now supports high woody-plant diversity, with 91 native species across 34 families. Aboveground biomass (AGB) averaged 66.91 ± 41.2 Mg ha⁻¹, comparable to seasonally dry tropical forests globally. Although native species were planted more recently and are shorter than non-natives, they contributed 23.86 ± 23.4 Mg ha⁻¹ to AGB and show potential for future increases in basal area. Given their comparable wood densities and capacity to attain similar heights, native species are predicted to sequester carbon at levels similar to non-natives in the long term. AGB was unrelated to species diversity. Overall, native TDEF species can achieve carbon storage while maintaining ecological integrity.

---

HUMAN-INDUCED climate change, caused by emissions of heat-trapping gases such as carbon dioxide and methane, is now widely recognized as having severely impacted natural and human systems globally^1^. As a consequence, stabilizing or removing atmospheric carbon is a global priority, and ecosystem restoration, afforestation and reforestation are seen as important contributors towards achieving this goal^1^. Indeed, it is partly for this reason that 2021-2030 has been declared the United Nations Decade on Ecosystem Restoration^2^. In addition, reforestation initiatives can generate significant income from carbon-trading schemes^3^. Beyond carbon, ecosystem restoration is recognized as necessary for biodiversity conservation, sustainable development, poverty alleviation and improved human well-being^2^.

However, a significant portion of historical and recently pledged afforestation and reforestation efforts have been directed towards planting a few species of fast-growing, often non-native, but economically-important tree species^4,5^. This is problematic because restored natural forests, which consist of multiple plant species, can potentially sequester more carbon than plantations^4,6,7^. The positive association between plant species and functional diversity and carbon storage is well-recognized, and attributable to mechanisms such as sampling effects (e.g., greater carbon stocks in more diverse stands due to the increased probability of inclusion of species that have denser wood or larger adult size), niche complementary, and facilitation^8–16^. Another mechanism contributing to overyielding is neighbourhood-mediated plasticity in biomass allocation and crown architecture in diverse stands^17^. Finally, positive plant diversity-carbon storage relationships have been demonstrated to strengthen over time due to trait-dependent shifts in overyielding species^18^. These mechanisms also apply to mixed species plantations, tending to make them more productive than monocultures^19^.

Just as importantly, restored natural forests can be superior at providing a range of beneficial ecosystem services such as biodiversity conservation, soil erosion control, and water provisioning, while also being more resilient to environmental change^7,20–22^. Furthermore, non-native trees favoured for plantations tend to be fast-growing and hence resource acquisitive, which can decrease their long-term carbon sequestration capacity due to lower carbon durability, despite being able to sequester carbon faster in the short-term^23^. In particular, in dry climates, native species can have greater carbon gain by recruitment and lower carbon loss by mortality than non-natives^23^. Thus, there is an urgent need to migrate from species-poor plantations, often consisting of non-native species, to natural, high-diversity, resilient ecosystems. Quantification of carbon stocks in non-native species plantations that are being restored to natural forests can provide important insights into the carbon sequestration potential of this pathway.

Tropical dry evergreen forests (TDEFs) are a unique forest type found in the seasonally-dry regions in the tropics of Central America, Africa, and Asia that are dominated by evergreen species. In Asia, these short-statured, closed-canopied forests are confined to Thailand, Sri Lanka, and India^24^. In India, TDEFs only occur within a relatively small area on the southeastern (Coromandel) seaboard of the Indian peninsula, ranging inland between 30 km^25^ and 60 km^26^. By a recent remote-sensing based estimate, TDEFs today occupy an area of 378 km^2^ in India, making up 0.06% of the total forest area and 0.01% of the total land area^27^. A few prior studies suggest TDEFs have considerable carbon sequestration potential and harbour high plant and animal diversity^24,28,29^. Southern Indian TDEFs have historically occurred in regions of high human population densities and have therefore largely been converted to other land-uses, only existing today in a few isolated patches^24,28,30^. Thus, southern Indian TDEFs have a high restoration potential. However, there is a paucity of data on the carbon sequestration potential of TDEFs, particularly those that have been restored. In particular, it is unclear how native TDEF species compare with the economically-favoured, faster-growing, non-native species in terms of carbon sequestration potential. Finally, it is unclear if biomass increases with species diversity in TDEF, as has been observed in other ecosystems.

The present paper addresses these gaps by examining a restored southern Indian TDEF. Located in Auroville, the site was originally largely barren in 1973 and was initially planted mainly with non-native tree species. The focus shifted to restoring TDEF in the early 1990s, and non-native species have steadily been replaced with native TDEF species ever since.

Our objectives were to:

a. quantify woody-plant diversity, stand structure, and carbon stocks,
b. quantify the individual contributions of native and non-native species to diversity, structure, and carbon stocks,
c. assess whether native TDEF species, which were planted more recently than non-natives, have comparable carbon sequestration potential in the future as non-natives, and
d. test the hypothesis that aboveground biomass (AGB) increases with species diversity.

## Materials and methods

### Study area

The Coromandel Coast receives about 1250 mm mean annual precipitation overall, though the amount varies greatly from north to south and follows a highly seasonal pattern with light rains from June to September and sporadic heavy falls from October to December, mostly resulting from depressions forming in the Bay of Bengal (the northeastern monsoon)^30,31^. TDEFs are associated with areas that face strong winds and/or have soils that drain easily, leaving soil moisture insufficient to balance evapotranspirative losses^32^. The soils of TDEFs are primarily ferrallitic sandy loam, which forms the “Cuddalore sandstone” of the Miocene period. This sandstone’s fossils contain indigenous floristic elements of southern India, which include >39% typical TDEF species but less than 10% companion (*Albizia amara* communities) and dry deciduous species^33^.

The study was carried out in Auroville, a town in southern India (Figure 1). The region has a dry season of about 6 months and receives rainfall mainly from the southwest (June to September) and northeast (November to December) monsoons. The mean annual rainfall was 1600 mm during 2019-2023. The mean maximum and minimum temperatures are 32.2 °C and 20 °C, respectively (adapted from Aurogreen Auroville Rainfall Data 2023).

**Figure 1.**
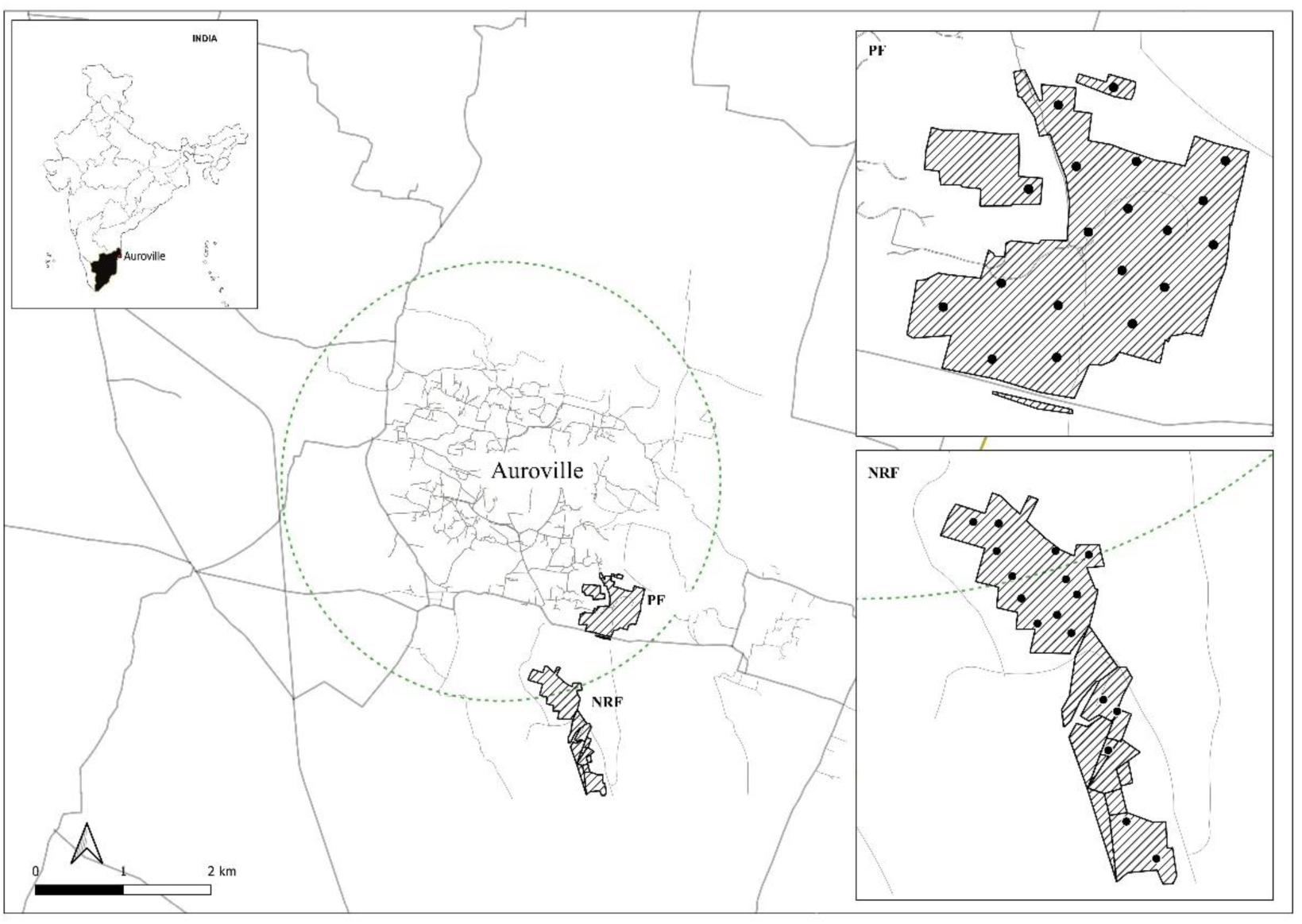
Map showing the location of the study site and sampling plots in Auroville, Tamil Nadu, India.

As a result of historical anthropogenic disturbances including logging and livestock grazing, prior to afforestation, the study site was largely barren, with a patchy cover of thorny shrubs and grasses, and dissected by deep gullies that carried off exposed soil into the sea during monsoons^30^. Afforestation was initiated in 1973 using green manure to rebuild the soil and by planting mainly non-native tree species including *Acacia auriculiformis*, *Acacia mangium*, *Acacia colei*, and *Eucalyptus globulus* to provide windbreaks and shade. Starting in the early 1990s, the focus shifted to TDEF restoration, which involved replacing non-native trees with native plants to restore ecosystem integrity, support biodiversity, and ensure long-term sustainability by favouring species better adapted to local conditions^34^. Native plant species subsequently started to recolonize and regenerate naturally. The total carbon stock in Auroville was estimated at 8605 Mg based on an analysis of LANDSAT TM data, with a tree cover estimate of 920 hectares between February 2017 and February 2022^35^. However, there are currently no ground-based studies on the carbon stocks or structure of Auroville’s restored forests.

### Tree inventory

Vegetation was sampled at two sites (i) Pitchandikulam Forest, a 30-hectare restored TDEF patch (11°59’39.83”N, 79°49’18.79”E; 73 m ASL; hereafter PF), and (ii) New Land-Ravena Forest, a 43- hectare restored TDEF patch (11°58’42.38”N, 79°49’6.43”E; 69 m ASL; hereafter NRF). The sites are similar, though compared to PF, NRF is in a lower-lying area that receives runoff from higher up in the watershed.

Trees were measured in 20 m × 20 m vegetation plots, placed randomly, spaced at least 80 m apart. 36 plots were laid in total: 19 in PF (0.76 ha) and 17 in NRF (0.68 ha). Steep terrain prevented the laying of square plots at three locations (1 in PF and 2 in NRF) and rectangular plots of the same area (10 m x 40 m) were laid instead. All trees with girth at breast height (GBH; measured 1.37 m from the ground) ≥ 10 cm were identified to the species and measured for diameter at breast height (DBH). Heights of trees taller than about 4 m were measured using laser rangefinder (BOSCH GLM 50-23G; Robert Bosch Power Tools Sdn Bhd, Malaysia) in conjunction with the smartphone app ‘Trees’ (Forest Monitoring Tools version 4.1.8). Heights of trees shorter than about 4 m were measured using a 4 m measuring pole. Fourteen tree heights had to be estimated by eye due to poor accessibility.

### Data analysis

Information on the native geographical range of each species was obtained from the Plants of the World Online Portal (https://powo.science.kew.org), supplemented by available literature on the species. *Pterocarpus santalinus* (Fabaceae), although naturalized in Tamil Nadu, is native to the Eastern Ghats in the neighbouring state of Andhra Pradesh and was therefore considered non-native in our analysis. In order to compare the diversity of native and non-native species and to assess the relationship between diversity and AGB, for each plot, three widely-used diversity measures were calculated: (a) species richness, (b) Shannon index, H’ = -∑ (*n_i_*/N) ln(*n_i_*/N), where N is total number of individuals, and *n_i_* is the number of *i*th species, and (c) Simpson index, D = 1 - ∑ (*n_i_*/N)^2^. Both Shannon and Simpson index take into account species abundances and represent both richness and evenness, while Simpson index is slightly more sensitive to the abundance of common species. We also calculated Hill numbers of order 1 (= exp(H’)) and order 2 (= 1/(1-D)). Hill numbers measure the effective numbers of species, i.e., the number of equally abundant species that would be needed to give the same diversity value. In order to characterize the relative dominance of species in this plant community, the Importance Value Index (IVI) of each species was calculated (IVI = relative density + relative frequency + relative dominance).

Following^36^, above ground biomass (AGB, Mg) was estimated using allometric equation AGB = 0.0673 × (ρD^2^H)^0.976^ × 0.001, where D is the DBH in cm, H is the height in m, and ρ is the wood density in g cm^-3^. This model has been shown to be the most accurate across forest types and bioclimatic conditions^36^. Wood density was obtained at the species-level from previously published studies^29,37^ (worldagroforestry.org/wd). Genus-level and family-level mean wood densities were used for 5 species each for which species-level wood density was unavailable. Aboveground carbon density, AGC (Mg C) was calculated as AGC = Σ_i_(AGB_i_ × CF_i_), where AGB_i_ is the AGB and CF_i_ is the carbon fraction (Mg C Mg^-1^ dry matter) of individual i. CF was set to 0.47 Mg C Mg^-1^ dry matter^38^ (V4, Ch4, Table 4.38). Belowground carbon (BGC) was calculated as BGC = AGC × 0.275^39^. Total carbon density (TC, Mg C) was calculated as TC = AGC + BGC.

AGB, and its constituents – density, basal area, height, and wood density – were compared between native and non-native species using Welch’s two-sample t-tests. Native species have been growing for a shorter duration at the site compared to non-native species and are therefore less likely to have achieved their maximum DBH potential. To compare future carbon storage potential, we compared how heights scale with DBH across native and non-native species. The height-DBH relationship was quantified using the linear regression log(height) ∼ log(DBH)*Origin, where ‘Origin’ indicated whether the individual was native or non-native. A steeper height-DBH relationship would indicate greater height, and therefore greater carbon storage, at a given DBH.

The relationship between plot-level AGB and each of the three diversity measures was assessed using linear regressions. All calculations were performed in R^40^. AGB was calculated using the package `BIOMASS`^41^.

## Results

1947 woody-plant individuals ≥ 10 cm GBH belonging to 117 species (34 families) were inventoried across thirty-six 400 m^2^ plots (total area 14,400 m^2^; Supplementary Figure 1). Of these, 1002 individuals belonging to 91 species (34 families) were native to the region, and 945 individuals belonging to 26 species (9 families) were non-native. Plot-level mean species- and family-richness was 16.3 and 9.5, respectively (Table 1). All diversity indices indicated higher mean plot-level native species diversity compared to non-native species diversity (Table 1). The non-native *Acacia auriculiformis* was the most abundant and dominant tree species by far (Table 2). Other dominants included *Khaya senegalensis*, *Libidibia ferrea* and *Pterocarpus santalinus*, all of which are non-native (Table 2; Supplementary Table 1).

**Table 1.** Means and standard deviations of plot-level estimates of measured variables for woody-plants censused in 36 vegetation plots. Diversity measures are reported as-is (i.e., at 400 m^2^ level); all other variables have been converted to a per-hectare level.

| Variable | All species | Native species | Non-native species |
| --- | --- | --- | --- |
| Density (ha<sup>-1</sup>) | 1352.08 ± 515.9 | 695.83 ± 379.5 | 656.25 ± 642.6 |
| Basal area (m<sup>2</sup> ha<sup>-1</sup>) | 19.49 ± 5.8 | 8.32 ± 6.6 | 11.17 ± 6.3 |
| Mean height (m) | 4.23 ± 0.9 | 3.48 ± 0.8 | 5.32 ± 1.3 |
| AGB (Mg ha<sup>-1</sup>) | 66.91 ± 41.2 | 23.86 ± 23.4 | 43.05 ± 36 |
| AGC (Mg ha<sup>-1</sup>) | 31.45 ± 20.2 | 11.47 ± 11.5 | 20.22 ± 17.6 |
| BGC (Mg ha<sup>-1</sup>) | 8.65 ± 5.6 | 3.08 ± 3.1 | 5.56 ± 4.9 |
| TC (Mg ha<sup>-1</sup>) | 40.09 ± 25.7 | 14.30 ± 14.6 | 25.79 ± 22.5 |
| Number of families | 9.5 ± 2.6 | 8.72 ± 2.8 | 1.92 ± 1 |
| Species richness | 16.28 ± 5.6 | 13.06 ± 5.5 | 3.22 ± 1.6 |
| Shannon index | 2.09 ± 0.7 | 2.23 ± 0.5 | 0.71 ± 0.5 |
| Simpson's index | 0.75 ± 0.2 | 0.85 ± 0.1 | 0.38 ± 0.3 |
| Hill number order 1 | 9.89 ± 5.6 | 10.23 ± 4.4 | 2.35 ± 1.4 |
| Hill number order 2 | 7.06 ± 4.9 | 8.32 ± 3.7 | 2.08 ± 1.3 |

**Table 2.**
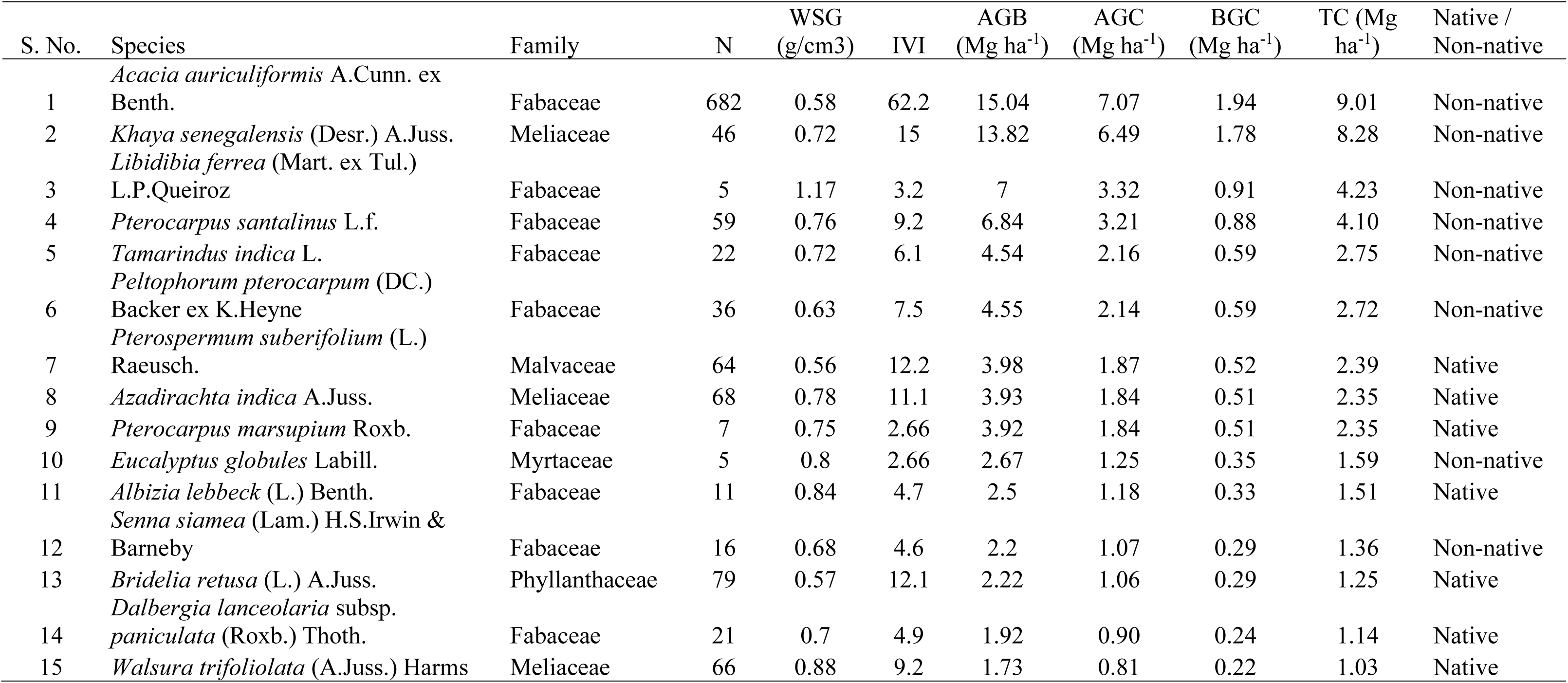
The top 15 woody-plant species censused in the 36 vegetation plots. Species are listed in descending order of total carbon (TC); N – abundance across all plots, WSG – wood specific gravity, IVI – Importance Value Index, AGB – aboveground biomass, AGC – aboveground carbon, BGC – below-ground carbon, TC – total carbon (= AGC + BGC). See Supplementary Table 1 for the complete list of species.

AGB in the plots was estimated at 66.91 ± 41.2 Mg ha^-1^ (mean ± SD; range 19.5 Mg ha^-1^ – 188.2 Mg ha^-1^). Total carbon (above- and below-ground) was estimated at 40.09 ± 25.7 Mg ha^-1^ (range 11.7 Mg ha^-1^ – 112.8 Mg ha^-1^). AGB distribution across DBH classes was bell-shaped for both native and non-native species (Figure 2A). In the smaller (< 20 cm DBH) size classes, both native and non-native species had comparable AGB. However, in all the larger (≥ 20 cm DBH) size classes, non-native species had nearly double the AGB of native species. The population structure was reversed J-shaped both for native and non-native species (Figure 2B). However, the latter had a greater proportion of individuals in the ≥ 20 cm DBH classes (Figure 2B).

**Figure 2.**
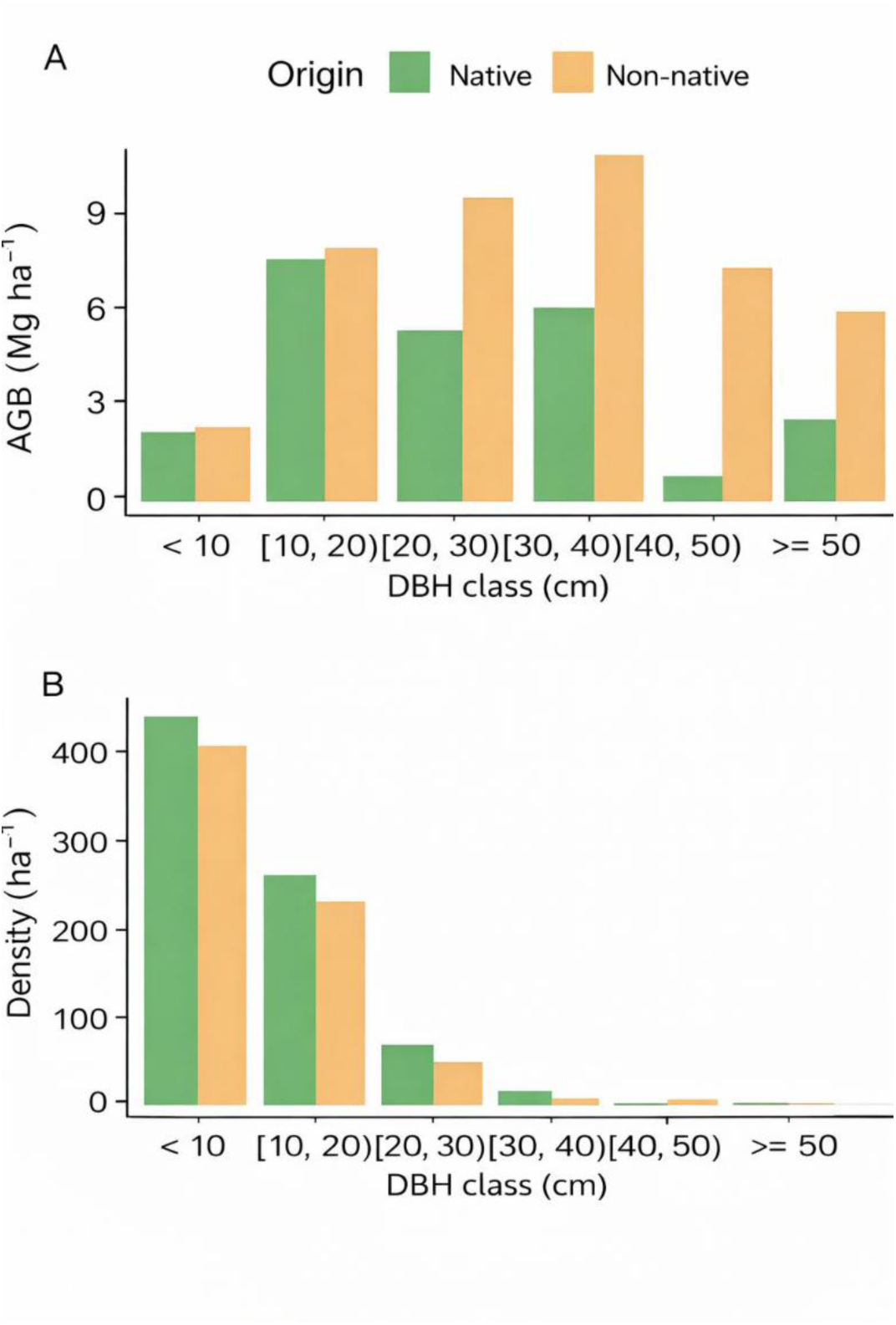
Aboveground biomass (AGB; Mg ha^-1^) (A: by species origin) and number of individuals (B: by species origin; ha^-1^) by DBH class (cm).

Non-native species showed no significant difference in abundance compared to native species, whether the dominant *Acacia auriculiformis* was included (Figure 3A; difference = - 17.58 ha^-1^, 95% CI [-54.85, 19.69], *P* = 0.341) or excluded (Figure 3B; difference = 0.34 ha^-1^, 95% CI [-4.83, 5.52], *P* = 0.895). Despite this similar abundance, non-native species possessed significantly greater ABG (Figure 3C; native - non-native difference = -1.39 Mg ha^-1^, 95% CI [-2.56, -0.23], *P* = 0.021). This biomass advantage was not driven by wood density, which remained comparable between the two groups (Figure 3D; difference = -0.01 g cm^-3^, 95% CI [-0.09, 0.07], *P* = 0.791). Instead, the greater AGB in non-native species was attributed to a marginally significant increase in basal area (Figure 3E; difference = -3382.37 cm^2^ ha^-1^, 95% CI [-7115.68, 350.94], *P* = 0.074) and, most critically, the fact that they were significantly taller (Figure 3F; difference = -2.9 m, 95% CI [-4.33, -1.47], *P* = 0.0002).

**Figure 3.**
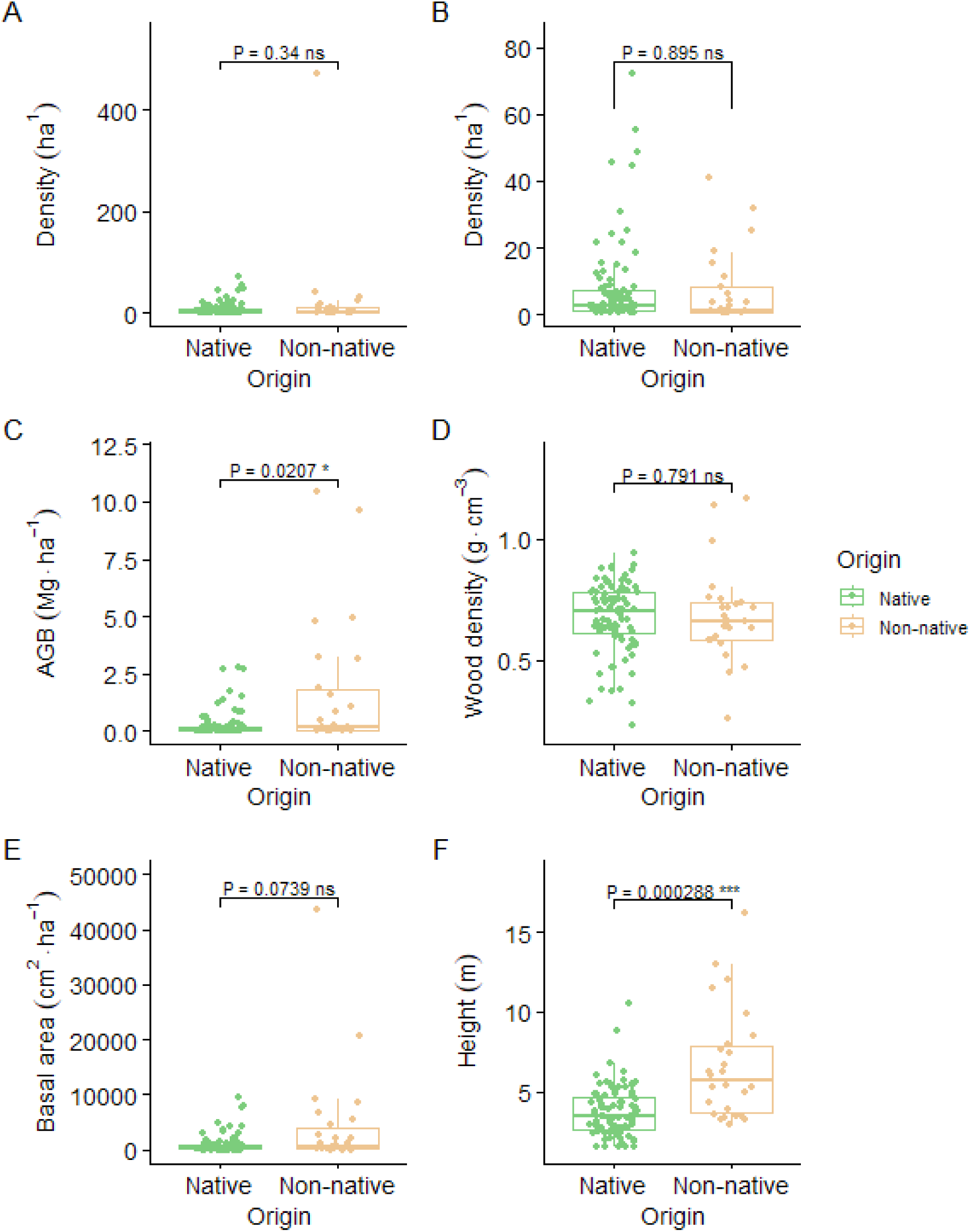
Comparisons between native and non-native species in terms of (A) number of individuals (ha^-1^), (B) number of individuals excluding the highly dominant non-native species *Acacia auriculiformis* (ha^-1^), (C) aboveground biomass (AGB; Mg ha^-1^), (D) wood density (g cm^-3^), (E) basal area (cm^2^ ha^-1^), and (F) mean height (m).

There was no significant relationship between AGB and either species richness (Figure 4A; slope = -0.11, 95% CI [-2.69, 2.46], *P* = 0.93), Shannon index (Figure 4B; slope = 9.3, 95% CI [-10.71, 29.31], *P* = 0.351), or Simpson index (figure not shown; slope = 35.75, 95% CI [-26.77, 98.27], *P* = 0.253). Similarly, there was no significant relationship between AGB and Hill numbers of order 1 and 2 (Figure S1).

**Figure 4.**
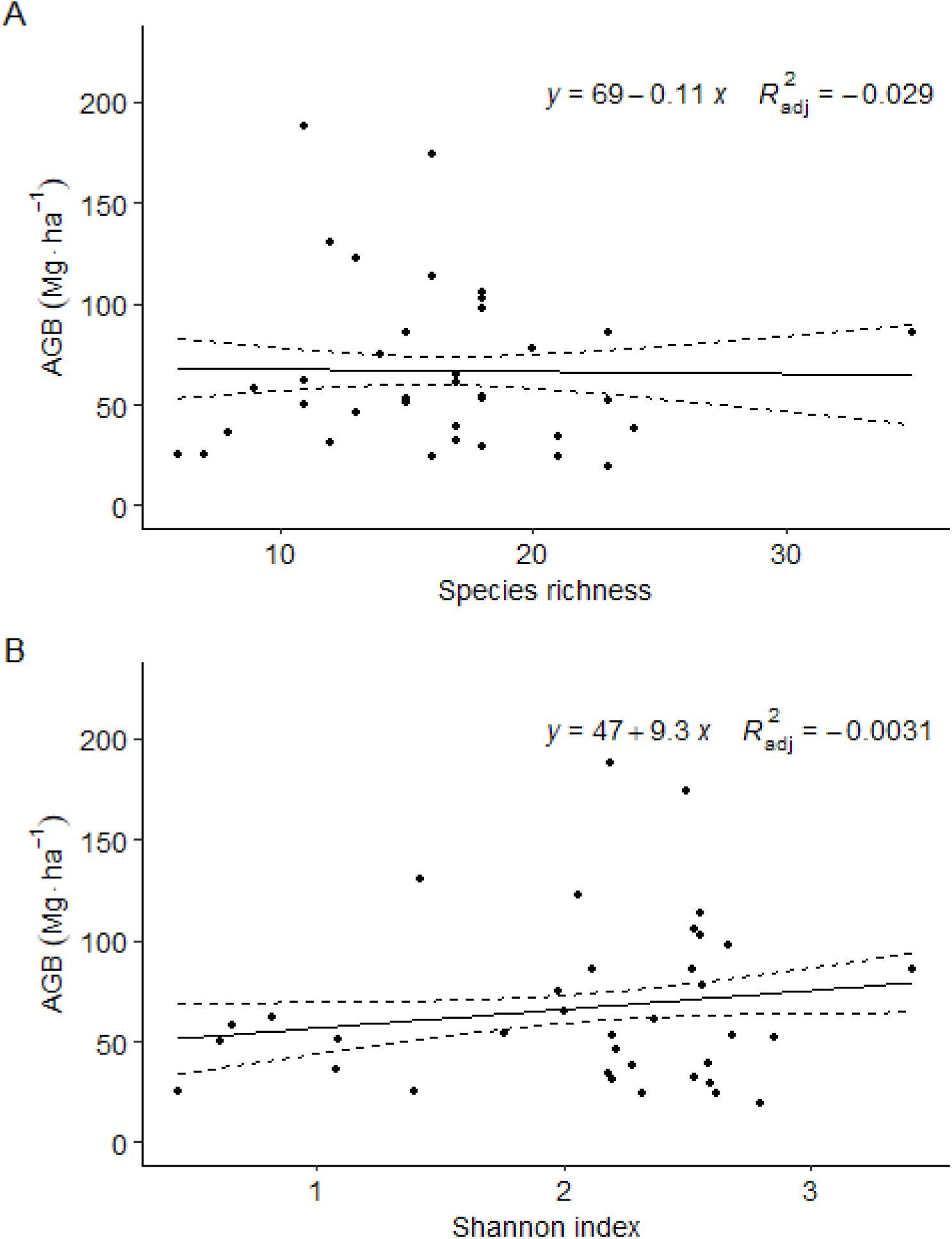
Linear regressions between aboveground biomass (AGB; Mg ha^-1^) and (A) species richness and (B) Shannon index.

Plant height (m) was strongly linearly related to DBH (cm) on log-log scale for both native (height = 0.81 × DBH^0.62^) and non-native species (height = 1.42 × DBH^0.5^) (Figure 5). The difference between native and non-native species in both the height-at-1-cm-DBH (0.623, 95% CI [0.588, 0.659], *P* < 0.001) and DBH exponent (-0.12, 95% CI [-0.17, -0.072], *P* = 0.000001) were significant. Thus, non-native species are significantly taller at small DBH values but this difference diminished with increasing DBH.

**Figure 5.**
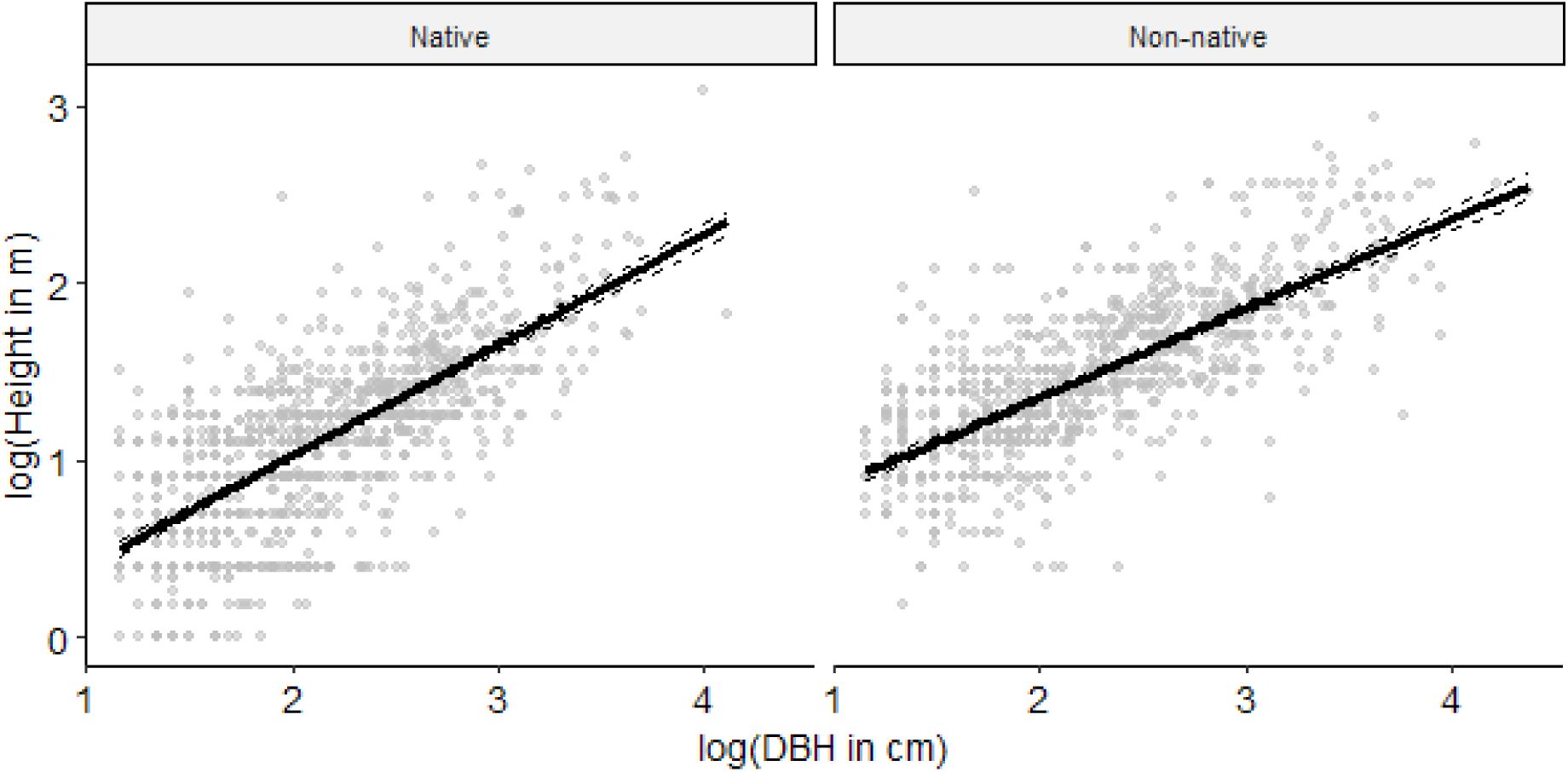
Linear regression between log-height (m) and log-DBH (cm) interacting with species origin (native vs. non-native).

## Discussion

Our study examined the diversity, structure, and carbon sequestration potential of a restored southern-Indian TDEF. Results from our analysis suggest that restoration of this threatened vegetation type can help achieve both high biodiversity (91 native woody-plant species belonging to 34 families at our site) and high biomass carbon density; the latter is comparable in magnitude to global seasonally dry tropical forests and several temperate and boreal forest types^42^. Carbon density estimates from the present study are consistent with previous estimates from TDEFs preserved within sacred groves in the region^29^. The mean AGB estimate from the present study is higher than the previous LANDSAT mean AGB estimate of 9.35 Mg ha^-1^ from Auroville^35^. This difference is likely due to two factors: (a) that study covered a larger area that includes less dense forests and, (b) unlike the present study, was based on AGB estimates from Forest Survey of India’s plot-based inventories of TDEFs elsewhere.

The reversed J-shaped size-class structure suggests either a stable or growing population, suggesting an even greater capacity to sequester carbon, though this needs to be confirmed by long-term monitoring^43^. Like in other forests, much of this carbon is stored in a relatively small number of large-sized individuals. For example, individuals with DBH > 20 cm contributed to 70.6% of AGB (68.1 Mg of 96.4 Mg) despite making up only 10.9% of individuals (213 of 1947). Large trees contribute disproportionately to stand biomass when compared to small trees because biomass grows exponentially with tree diameter^12^.

In this forest, non-native species dominated AGB and to a lesser extent density in the larger size classes, which reflects the fact that early planting efforts mostly involved non-natives. Comparison to natural regional TDEFs suggests that the native species basal area of 8.32 m^2^ ha^-1^ (of a total basal area of 19.49 m^2^ ha^-1^) can potentially increase as native species continue to grow (e.g., Baithalu et al.^44^ reported native species basal area of 37.5 m^2^ ha^-1^ for similar tree densities). The height-DBH models suggest that if native species continue to increase in DBH – which is expected as they have been planted more recently – their heights will be comparable to that of non-native species. Together with the fact that native species have comparable wood densities, this suggests that native TDEF species are capable to sequestering as much carbon as large non-native, commercially-favoured species such as *Acacia auriculiformis*. Thus, as older non-native woody-plants die and native woody-plants grow larger, carbon stocks in this forest will increasingly transition to the latter and total carbon may surpass current values. The results from Costa Rica suggest that while lower-yielding but stable native species may survive better in challenging conditions, high-yielding but unstable invasive species contribute to inconsistent plantation success^45^.

We found no evidence that greater woody-plant diversity was associated with AGB overyielding. This is contrary to theory^46^ and findings from numerous forests globally where a positive relationship has been observed between AGB and species richness at small spatial scales, though not at large spatial scales^10,12,14^. Diversity-AGB relationships are scale-dependent possibly because underlying mechanisms such as sampling effects, niche complementarity, and facilitation dominate at smaller spatial scales while environmental gradients dominate at larger spatial scales^10^. However, our data have a grain size of 400 m^2^, which is in the range of scales at which a positive AGB-diversity relationships have been widely reported. Possible reasons for the absence of a relationship at our site include (a) overyielding being weaker in TDEFs, thereby requiring a larger sample size in order to be detectable, and (b) overyielding being weaker in forests that are being restored, compared to old-growth forests. In particular, the dominance of *Acacia auriculiformis* in our sites in may be a potential confounding factor since it decreases plot-level diversity while simultaneously increasing AGB. In addition, relationships between species richness and ecosystem functioning can result from the identity of species rather by richness per se^47^, which can lead to idiosyncratic relationships in some situations. Future studies are needed to test whether the widely-reported AGB-diversity relationship applies to restored TDEFs as well.

## Conclusion

Fast-growing tree plantations are often seen as important to carbon sequestration^48–50^. For example, *Acacia mangium* plantations have been considered a Clean Development Mechanism (CDM) for climate change mitigation^51,52^. However, restoring native TDEF species in its ecoregion can potentially help achieve similar levels of carbon sequestration in the long term. Although not studied here, previous research suggests that TDEFs provide numerous ecosystem services such as traditional medicinal plants, enhanced aquifer recharge, food security, genetic resources, pollinator services, aesthetic, spiritual and cultural services, and a habitat for native fauna and funga^28^. An additional advantage of preferring native species is that ecosystem resilience tends to increase with biodiversity^53^ and good potential for large-scale reforestation^54^. Restored TDEF patches can also provide a genetic diversity safety net for a large number of species, giving conservationists time to develop plans for preventing the extinction of the relevant species^55,56^. The absence of a diversity-AGB relationship in these restored forests merits further investigation.

## ACKNOWLEDGEMENTS

This project was funded by the Project Coordination Group (PCG), Auroville Foundation. We are grateful to Joss Brooks, Jaap den Hollander and Regina Braun for site permissions, Island Lescure for laser rangefinder, Aurosylle Bystrom, M. Azhagappan, P. Karunakaran, P. Iyyappan, R. Giridharan, D. Sripathy and R. Selvam for logistics support and fieldwork.

## Supplementary Material

**Figure S1.**
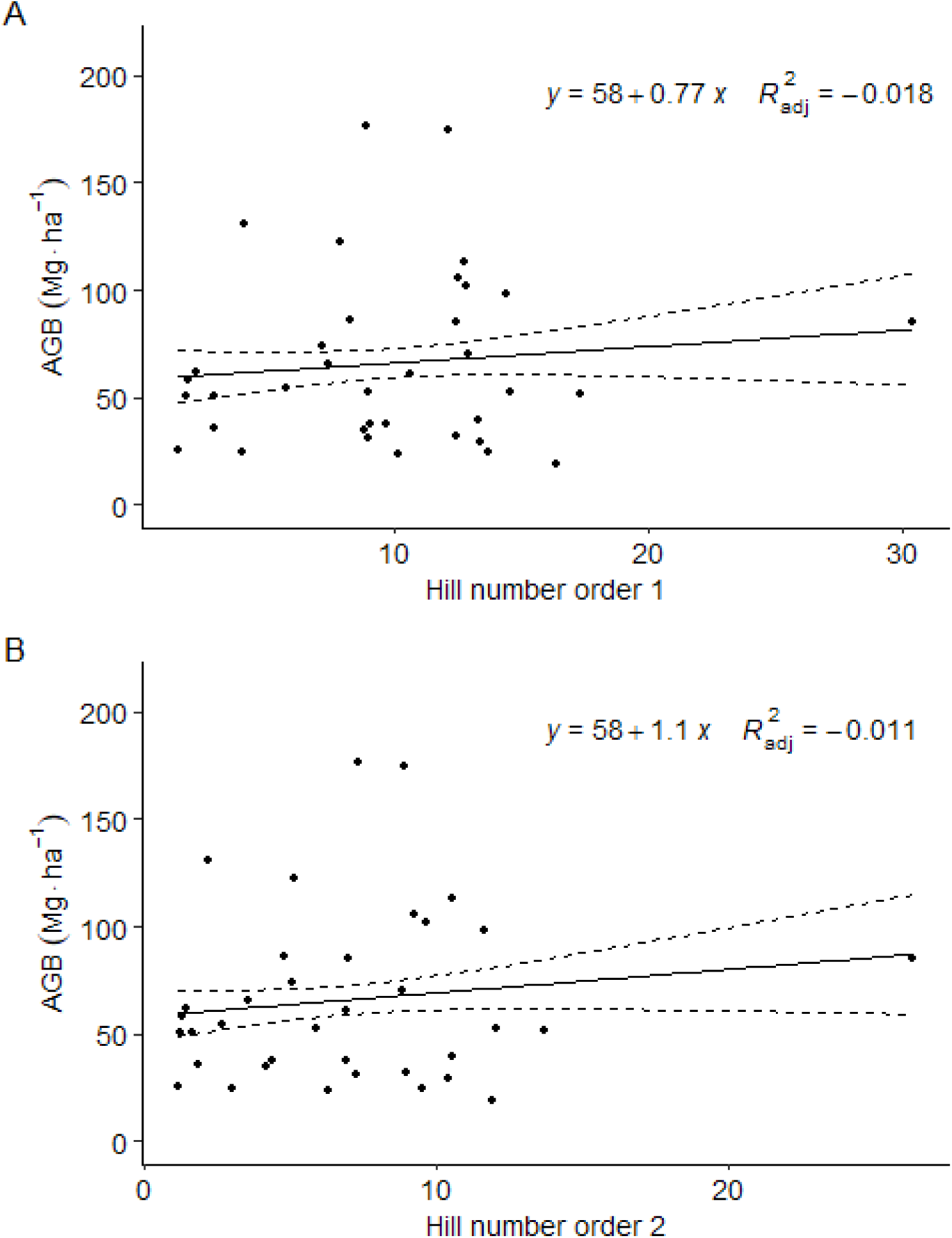
Linear regressions between aboveground biomass (AGB; Mg ha^-1^) and Hill numbers (A) of order 1 and (B) of order 2.

## Appendix 1

See Table 2

**Table 2.**
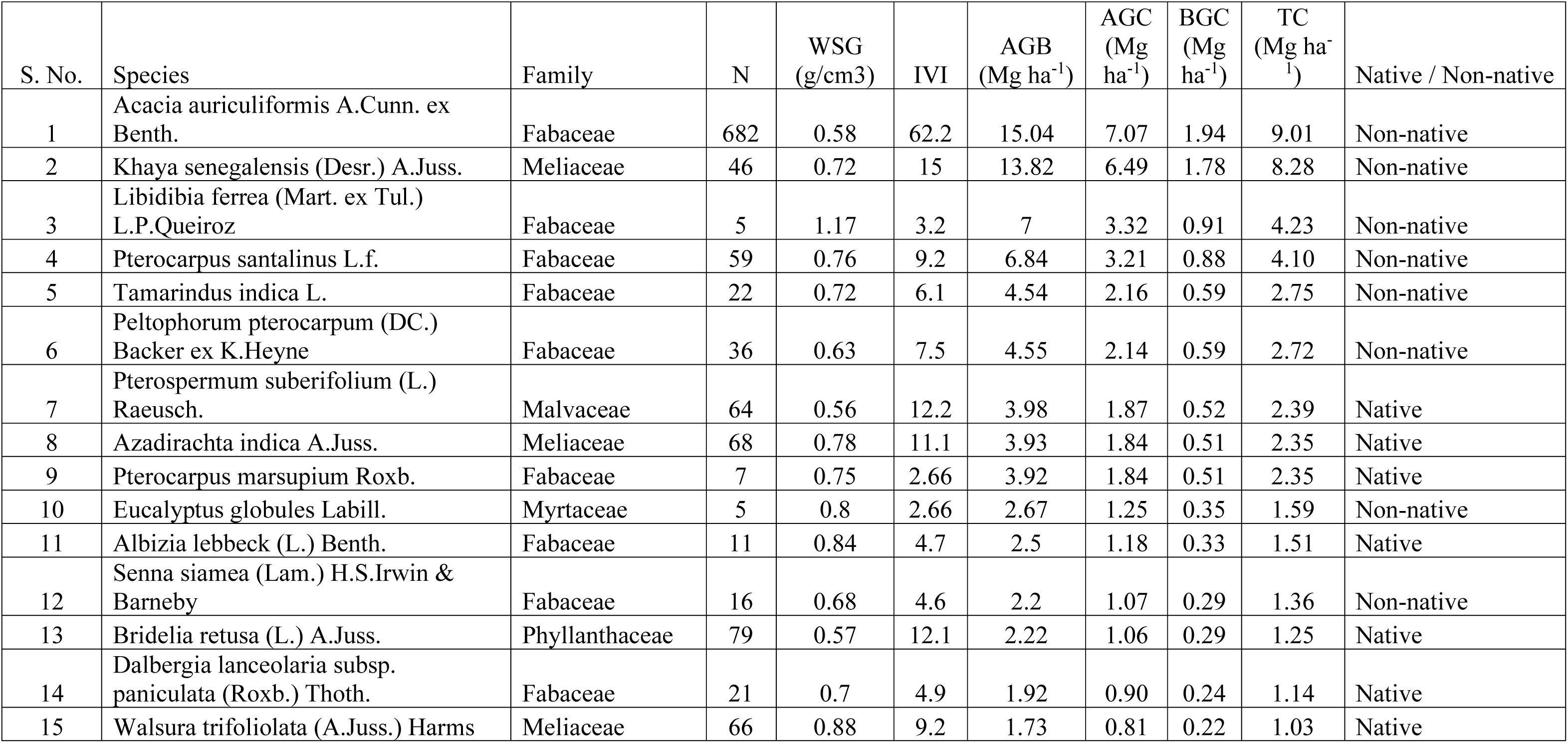
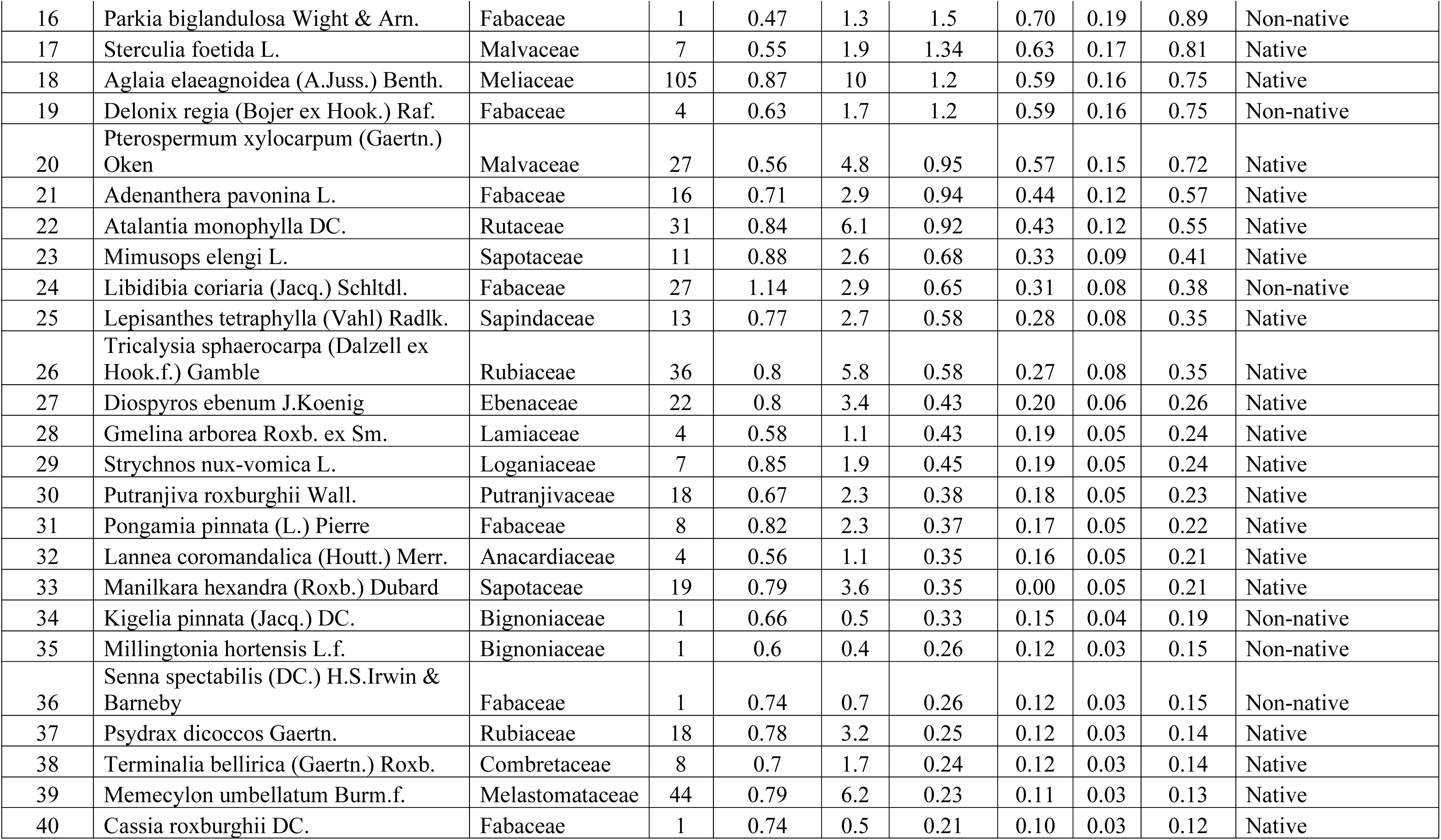
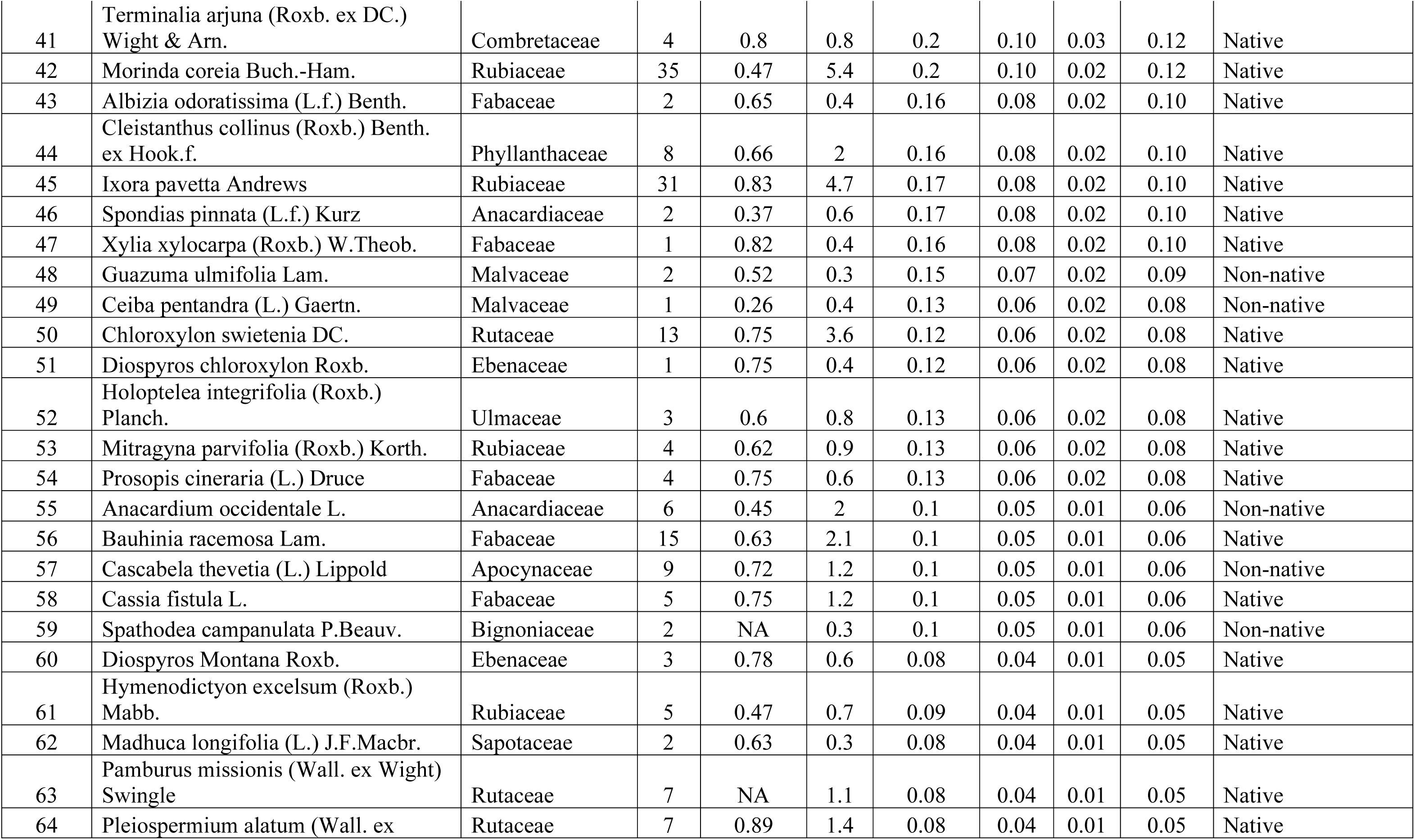
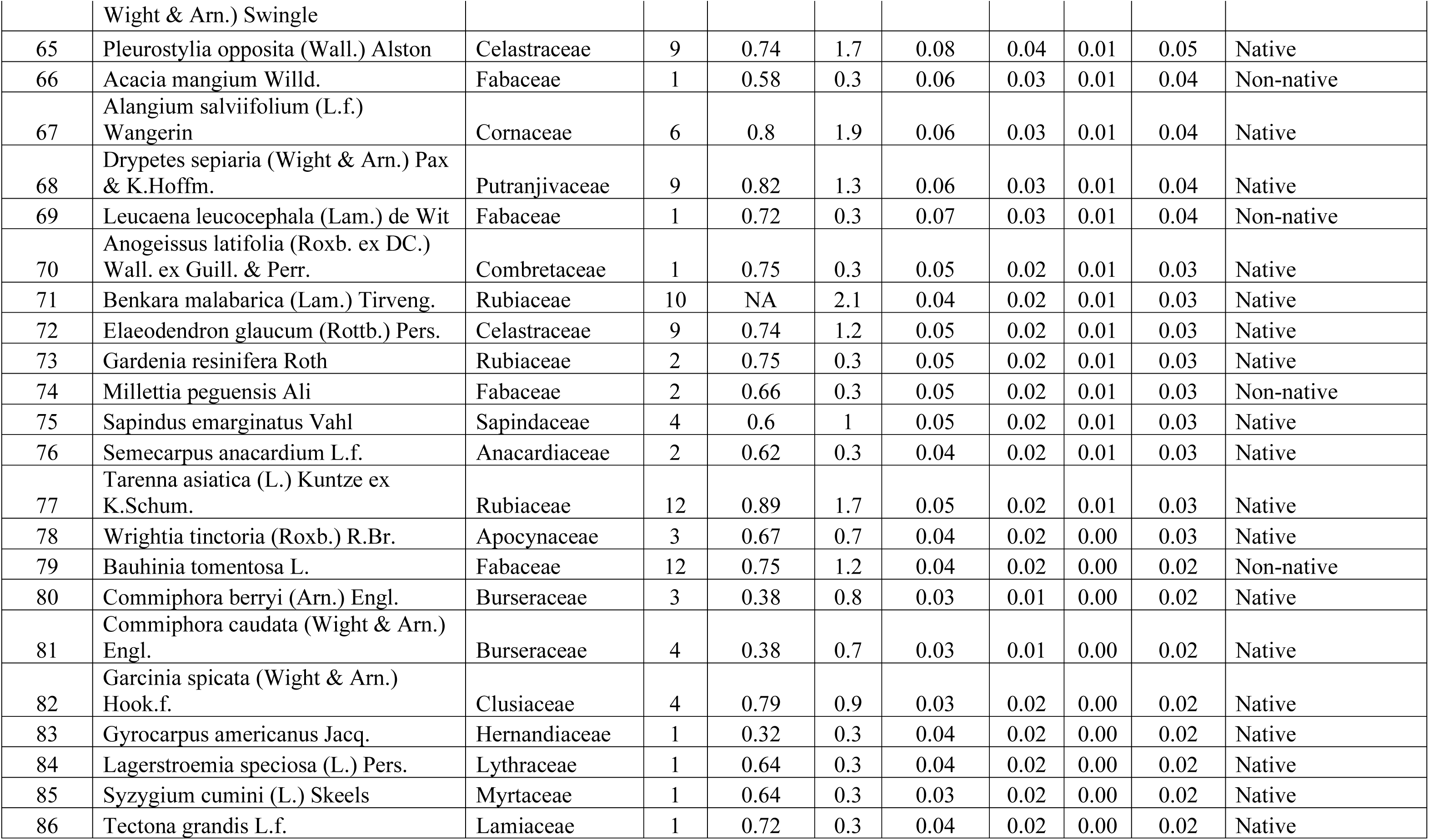
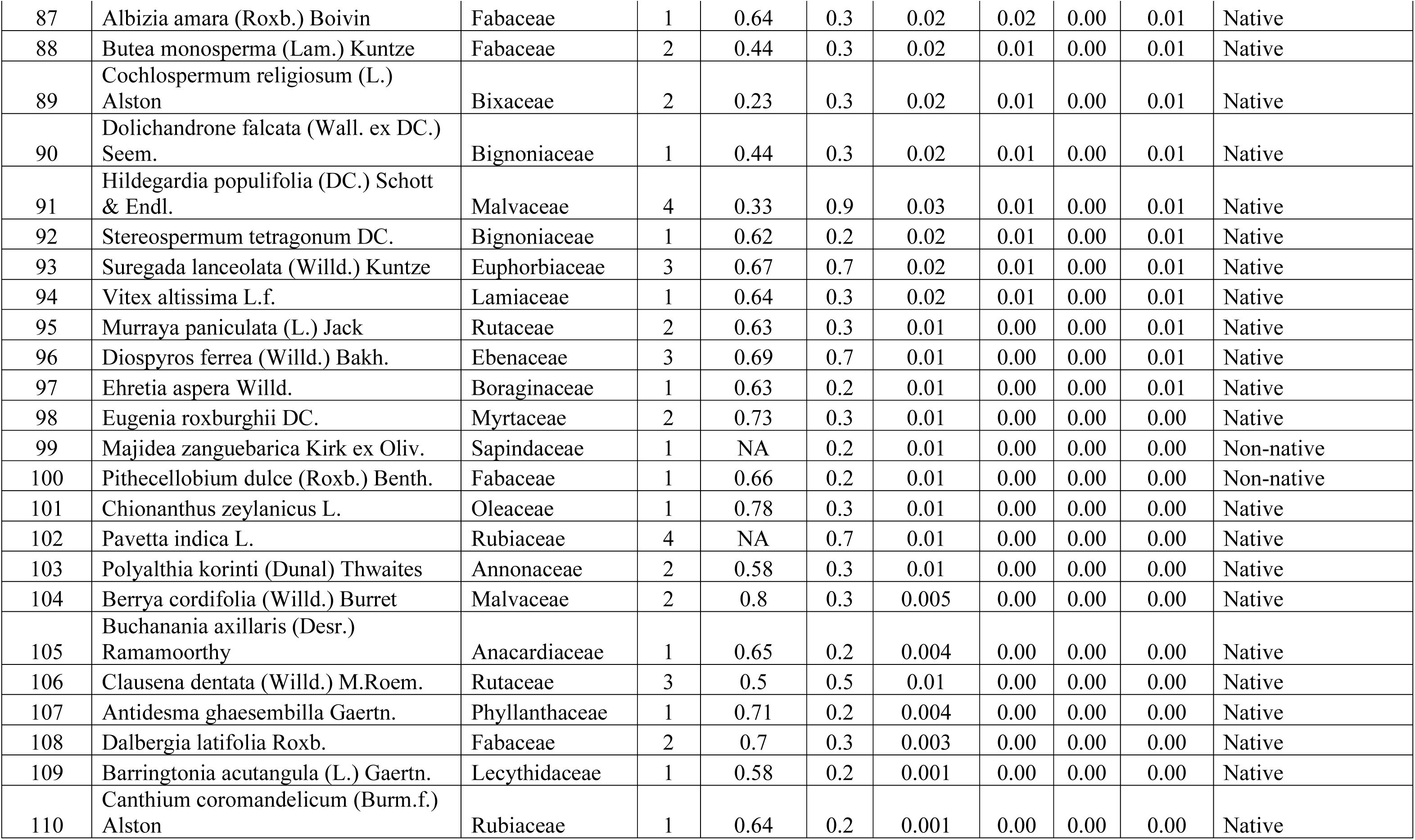
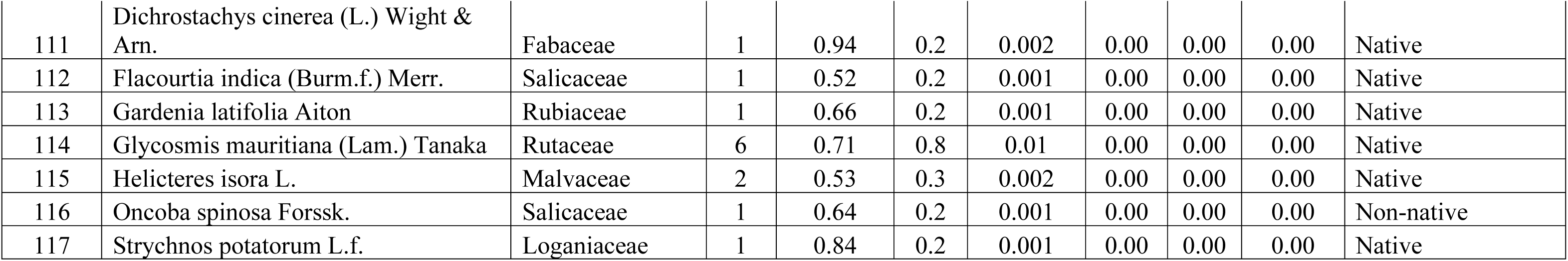
Woody-plant species censused in the 36 vegetation plots. Species are listed in descending order of total carbon (TC); N – abundance across all plots, WSG – wood specific gravity, IVI – Importance Value Index, AGB – aboveground biomass, AGC – aboveground carbon, BGC – below-ground carbon, TC – total carbon (= AGC + BGC), NA – not available.

## Appendix 2

See Figure 1

**Fig. 1.**
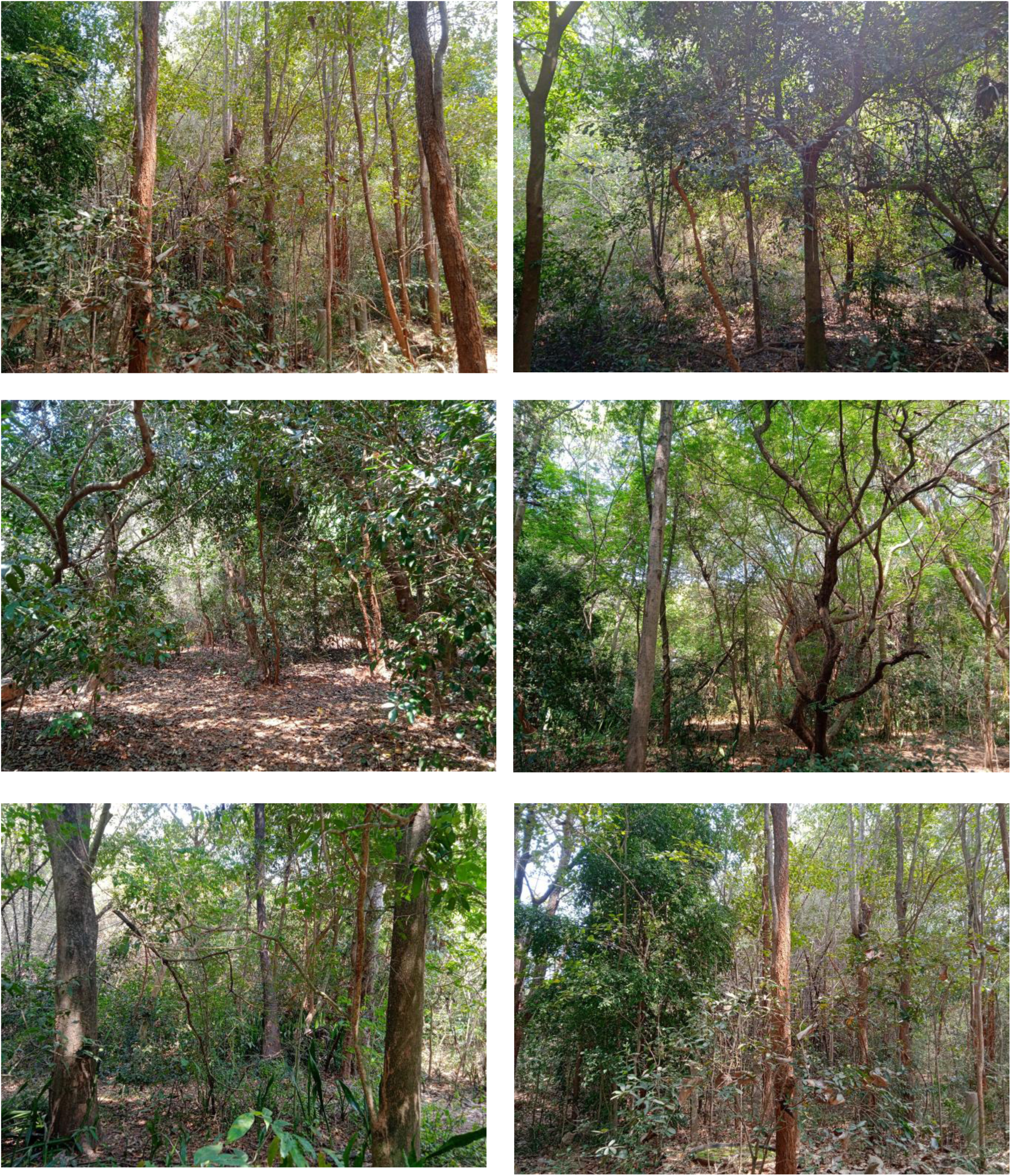
Images showing the structure of restored TDEF at various sampled locations

